# Quantitative consequences of protein carriers in immunopeptidomics and tyrosine phosphorylation MS^2^ analyses

**DOI:** 10.1101/2021.02.26.433124

**Authors:** Lauren E. Stopfer, Jason E. Conage-Pough, Forest M. White

## Abstract

Utilizing a protein carrier in combination with isobaric labeling to “boost” the signal of other low-level samples in multiplexed analyses has emerged as an attractive strategy to enhance data quantity while minimizing protein input in mass spectrometry analyses. Recent applications of this approach include pMHC profiling and tyrosine phosphoproteomics, two applications that are often limited by large sample requirements. While including a protein carrier has been shown to increase the number of identifiable peptides in both applications, the impact of a protein carrier on quantitative accuracy remains to be thoroughly explored, particularly in relevant biological contexts where samples exhibit dynamic changes in abundance across peptides. Here, we describe two sets of analyses comparing MS2-based quantitation using a 20x protein carrier in pMHC analyses and a high (∼100x) and low (∼9x) protein carrier in pTyr analyses, using CDK4/6 inhibitors and EGF stimulation to drive dynamic changes in the immunopeptidome and phosphoproteome, respectively. In both applications, inclusion of a protein carrier resulted in an increased number of MHC peptide or phosphopeptide identifications, as expected. At the same time, quantitative accuracy was adversely affected by the presence of the protein carrier, altering interpretation of the underlying biological response to perturbation. Moreover, for tyrosine phosphoproteomics, the presence of high levels of protein carrier led to a large number of missing values for endogenous phosphopeptides, leading to fewer quantifiable peptides relative to the “no-boost” condition. These data highlight the unique limitations and future experimental considerations for both analysis types and provide a framework for assessing quantitative accuracy in protein carrier experiments moving forward.

## INTRODUCTION

Mass spectrometry (MS)-based proteomics has historically been limited to analyzing bulk cell populations, largely due to losses during sample processing and limited instrument sensitivity. In recent years, several platforms have achieved protein expression profiling in single cells (e.g., single-cell proteomics (SCP)), a notable advancement in proteomics. To overcome sensitivity limitations and acquire deep proteomics datasets, the majority of these platforms rely on isobaric labeling (*i*.*e*., Tandem Mass Tags (TMT)) for sample multiplexing and a signal “boosting” sample, or “carrier proteome.”^1–3^ Carrier proteomes that have been utilized thus far contain a larger amount of protein than the non-carrier samples, an equivalent amount of protein but with a perturbation to increase the signal of interest^4^, or both.^5^ Because all isobaric labels have an identical intact mass, the inclusion of a carrier proteome increases the precursor ion intensity, enabling enhanced detection of low-input or low-level samples.

Use of a carrier proteome has also recently been applied to peptide major histocompatibility complex (pMHC) profiling (e.g., immunopeptidomics), and tyrosine phosphorylation (pTyr) analyses, both of which historically have required large sample inputs for sufficient signal detection by MS. For example, recent advances in pMHC profiling methods have decreased sample input requirements from >10^9^ cells to ∼10^7^ cells, yet even this lower boundary still represents a major limitation in the clinical translatability of the approach.^6,7^ Clinical specimens, including fine needle biopsies, typically do not provide enough material for deep pMHC profiling, and neoantigens are challenging to identify by MS, even with large sample quantities.^8^ Similarly, profiling pTyr peptides is possible using several hundred micrograms of input protein per channel in a multiplexed analysis^9^, but there is continued effort to reduce sample requirements to enable pTyr profiling of fine needle biopsies, tissue sections, or even single cells.

Inclusion of a protein carrier has resulted in an increased number of identifiable peptides in multiplexed immunopeptidomics analyses as well as multiplexed phosphotyrosine analyses. Ramarathinam et al. utilized increased protein material, cellular or patient-derived xenograft tumors, as a protein carrier in class I pMHC experiments, while Chua et al. used a protein carrier that had been treated with pervanadate (PV) treatment to halt tyrosine phosphatase activity and thereby increase tyrosine phosphorylation levels.^5^. While these initial results are encouraging, the quantitative impact of boosting in both approaches remains poorly understood. Specifically, a carrier proteome may limit the instrument’s dynamic range, leading to reporter ion ratio compression, and increase the number of missing values, thereby reducing data quality and/or data quantity, potentially altering biological interpretation.^10^

Several studies have begun to address these critical questions, albeit with limitations. For instance, experiments to assess ratio suppression typically evaluate whether constant ratios of protein input material are preserved in the presence of a protein carrier, which is not reflective of many biological systems where subtler changes in a subset of peptides demonstrate altered quantitation.^5,10^ Studies have also evaluated whether principal component analysis (PCA) can resolve differences between two cell populations in the presence of various protein carrier-to-signal ratios. However, these experiments generally use distinct cell types or cell lines, which have higher heterogeneity in peptide quantitation.^1,5,10,11^

Here, we describe results from analyses comparing MS^2^-based quantitation with and without the inclusion of a 20x protein carrier in pMHC analyses and a high (∼100x) or low (∼9x) carrier in pTyr analyses. We utilized isotopically labeled pMHCs to estimate changes in ion suppression in pMHC analyses along with cyclin-dependent kinase 4/6 (CDK4/6) inhibitor treatment to shift the pMHC repertoire in pathways related to cell cycle control.^7^ in pTyr experiments, epidermal growth factor (EGF) stimulation was used to drive a temporal pTyr response in a subset of the tyrosine phosphoproteome.^12^ In both applications, protein carriers altered peptide quantitation compared to the control experiment, inhibiting our ability to accurately interpret the biology underlying the cellular perturbations. Using these data, we define existing limitations for MS^2^-based analyses using protein carriers and highlight areas for future exploration that may enhance data quality through altered experimental design or acquisition framework.

## EXPERIMENTAL PROCEDURES

### Cell lines

SKMEL5 and A549 cell lines were obtained from ATCC [ATCC HTB-70 and CCL-185, respectively). Cells were maintained in DMEM medium (Corning) supplemented with 10% FBS (Gibco) and 1% penicillin/streptomycin (Gibco). Cells were routinely tested for mycoplasma contamination and maintained in 37°C, 5% CO_2_. Experiments were performed on passages 4-8.

### UV-mediated peptide exchange for hipMHCs

UV-mediated peptide exchange was performed using recombinant, biotinylated Flex-T HLA-A*02:01 monomers (BioLegend), using a modified version of the commercial protocol as previously described.^7^ Concentration of stable complexes following peptide exchange was quantified using the Flex-T HLA class I ELISA assay (BioLegend) as per the manufacturer’s instructions. ELISA results were acquired using a Tecan plate reader Infinite 200 with Tecan icontrol version 1.7.1.12.

### Synthetic peptide standards

Heavy leucine-containing peptides were synthesized at the MIT Biopolymers and Proteomics Lab using standard Fmoc chemistry using an Intavis model MultiPep peptide synthesizer with HATU activation and 5 μmol chemistry cycles. Starting resin used was Fmoc-Amide Resin (Applied Biosystems). Cleavage from resin and simultaneous amino acid side chain deprotection was accomplished using: trifluoroacetic acid (81.5% v/v); phenol (5% v/v); water (5% v/v); thioanisole (5% v/v); 1,2-ethanedithiol (2.5% v/v); 1% triisopropylsilane for 1.5 hours. Fmoc-Leu (^13^C_6_, ^15^N) was obtained from Cambridge Isotope Laboratories, and standard Fmoc amino acids were from NovaBiochem.

Peptides were subjected to quality control by mass spectrometry and reverse phase chromatography using a Bruker MiroFlex MALDI-TOF and Agilent model 1100 HPLC system with a Vydac C18 column [300 angstrom, 5 micron, 2.1 x 150 mm] at 300 μL/min monitoring at 210 and 280 nm with a trifluoroacetic acid/ H_2_O/MeCN mobile phase survey gradient.

### Peptide MHC isolation & TMT labeling

Cells were seeded in 10 cm plates and treated the following day for 72 hours with DMSO control, palbociclib (Selleckchem, PD-0332991), 10 ng mL^-1^ human recombinant IFN-γ (ProSpec Bio). During harvest, cells were washed with 1x PBS, and lifted with 0.05% Trypsin-EDTA (Gibco). Cells were pelleted, washed with 1x PBS, pelleted again, and resuspended in lysis buffer [20 nM Tris-HCl pH 8.0, 150 mM NaCl, 0.2 mM PMSF, 1% CHAPS, and 1x HALT Protease/Phosphatase Inhibitor Cocktail (Thermo Fisher)], followed by brief sonication to disrupt cell membranes. Lysate was cleared by centrifugation and quantified using bicinchoninic acid protein assay kit (Pierce).

Peptide MHCs were isolated by immunoprecipitation (IP) as previously described.^7^ Briefly, using 100 μg of pan-specific anti-human MHC Class I (HLA-A, HLA-B, HLA-C) antibody (clone W6/32, Bio X Cell) per 1e^6^ cells, which was bound to 10 μL FastFlow Protein A Sepharose bead slurry (GE Healthcare) per 1e^6^ cells for 3 hours rotating at 4°C. Beads were washed 2x with IP buffer (20 nM Tris-HCl pH 8.0, 150 mM NaCl), after which lysate/hipMHCs were added and incubated rotating overnight at 4°C. Beads were washed with 1x TBS and water, and pMHCs eluted in 10% formic acid for 20 mins at RT. Peptides were isolated from antibody and MHC molecules using a passivated 10K molecule weight cutoff filters (PALL Life Science), lyophilized, and stored at -80°C prior to TMT labeling.

To label pMHCs, 50 μg of pre-aliquoted Tandem Mass Tag 6-plex (TMT-6, Thermo Scientific) was resuspended in 20 μL anhydrous acetonitrile, and lyophilized peptides were resuspended in 66 μL 150 mM triethylammonium bicarbonate, 50% ethanol. TMT/peptide mixtures were incubated on a shaker for 1 hour at RT followed by 15 mins of vacuum centrifugation. Samples were next combined and centrifuged to dryness. Sample cleanup was subsequently performed using SP3, as previously described.^7,13^

### pTyr sample preparation

A549 cells were seeded in 10 cm plates and serum depleted for 72 hours prior to analysis. In EGF stimulation experiments, cells were stimulated with 5 EGF (PeproTech), flash frozen in liquid nitrogen, and lysed in 8M urea. Pervanadate treated cells were incubated for 30 mins with 30 µM pervanadate at 37°C prepared using 200 mM sodium orthovanadate, 1X PBS, and 30% hydrogen peroxide, followed by a 15 min incubation at RT protected from light. Cells were subsequently washed 1X with ice cold 1X PBS and lysed in 8M urea.

Lysates were cleared by centrifugation at 5000 g for 5 min at 4°C and protein concentration was measured by BCA (Pierce). Proteins were reduced with 10 mM DTT for 30 min at 56°C, alkylated with 55 mM iodoacetamide for 45 min at RT protected from light, and diluted 4-fold with 100 mM ammonium acetate, pH 8.9. Proteins were digested with sequencing grade modified trypsin (Promega) at an enzyme to substrate ratio of 1:50 overnight at RT. Enzymatic activity was quenched by acidifying with glacial acetic acid to 10% of the final solution volume, and peptides were desalted using C18 solid phase extraction cartridges (Sep-Pak Plus Short, Waters). Peptides were eluted with aqueous 40% acetonitrile in 0.1% acetic acid and dried using vacuum centrifugation. Peptide concentration was measured by BCA to account for variation in sample processing, and peptides were subsequently lyophilized.

Lyophilized peptides were labeled with TMT-10plex in ∼35 mM HEPES and ∼30% acetonitrile at pH 8.5 for 1 hour at room temperature. 100 µg peptide aliquots utilized 400 µg TMT, 900 µg-1 mg aliquots used 1600 µg TMT. Labeling reactions were quenched with 0.3% of hydroxylamine, and samples were pooled, dried in vacuum centrifugation, and stored at -80°C prior to analysis.

Labeled peptide aliquots were resuspended in 400 μL of immunoprecipitation (IP) buffer [100 mM Tris-HCl, 0.3% NP-40, pH 7.4] and incubated with 60 μL protein G agarose bead slurry (Calbiochem) conjugated to an antibody cocktail containing 24 μg 4G10 (Millipore) and 12 μg PT66 (Sigma), rotating overnight at 4°C. Beads were washed 1x with IP buffer, 3x with 100 mM Tri-HCl, pH 7.4, and eluted in 2 rounds of 25 μL 0.2% TFA. Phosphopeptides were further enriched using High-Select Fe-NTA Phosphopeptide Enrichment Kit (Thermo Scientific) following manufacturer’s instructions with minor adjustments as previously described.^14^ Peptide elutions were dried down using vacuum centrifugation to <2 μL total volume and resuspended in 5% acetonitrile in 0.1% formic acid for a total volume of 10 μL.

### MHC MS data acquisition

pMHC samples were analyzed using an Orbitrap Exploris 480 mass spectrometer (Thermo Scientific) coupled with an UltiMate 3000 RSLC Nano LC system (Dionex), Nanospray Flex ion source (Thermo Scientific), and column oven heater (Sonation). Samples were resuspended in 0.1% formic acid and directly loaded onto a 10-15 cm analytical capillary chromatography column with an integrated electrospray tip (∼1 μm orifice), prepared and packed in house (50 μm ID × 20 cm & 1.9 μM C18 beads, ReproSil-Pur). Twenty-five percent of pMHC elutions were injected for each analysis. Peptides were eluted using a gradient with 8-25% buffer B (70% Acetonitrile, 0.1% formic acid) for 50 mins, 25-35% for 25 mins, 35-55% for 5 mins, 55-100% for 2 mins, hold for 1 mins, and 100% to 3% for 2 mins.

Standard mass spectrometry parameters were as follows: spray voltage, 2.0 kV; no sheath or auxiliary gas flow; heated capillary temperature, 275 °C. The Exploris was operated in DDA mode. Full scan mass spectra (350-1200 m/z, 60,000 resolution) were detected in the orbitrap analyzer after accumulation of 3e^6^ ions (normalized AGC target of 300%) or 25 ms. For every full scan, MS^2^ were collected during a 3 second cycle time. Ions were isolated (0.4 m/z isolation width) for a maximum of 150 ms or 75% AGC target and fragmented by HCD with 32% nCE at a resolution of 45,000. Charge states < 2 and > 4 were excluded, and precursors were excluded from selection for 30 secs if fragmented n=2 times within 20 second window.

### pTyr MS data acquisition

LC-MS/MS analysis of pTyr peptides were performed on an Agilent 1260 HPLC system coupled to an Orbitrap Exploris 480 mass spectrometer. Peptides were resuspended in 10 μL 0.1% acetic acid and loaded onto an analytical capillary column with an integrated electrospray tip (∼1 μm orifice) prepared in house ((50 μm ID × 12 cm with 5 μm C18 beads (YMC gel, ODS-AQ, 12 nm, S-5 μm, AQ12S05)). Peptides were eluted using a 140-min gradient with 13-42% buffer B (70% Acetonitrile, 0.2M acetic acid) from 10-105 mins and 42-60% buffer B from 105-115 mins, 60-100% B from 115-122 mins, and 100-0% B from 128-130 mins at a flow rate of 0.2 mL/min for a flow split of approximately 10,000:1.

Standard mass spectrometry parameters were as follows: spray voltage, 2.5 kV; no sheath or auxiliary gas flow; heated capillary temperature, 275°C.

The mass spectrometer was operated in data-dependent acquisition with following settings for MS1 scans: m/z range: 350-2000; resolution: 60,000; AGC target: 3e^6^; auto IT: 50 ms. Within a 3 second cycle time, ions were isolated (0.4 m/z) and fragmented by HCD (nCE: 33%) with resolution: 60,000; AGC target: 1e^5^, max IT: 250 ms for all analyses except EGF-boost 500 ms (AGC target: 5e^5^, max IT: 500 ms). Unassigned and charge states <+2 and >+6 were excluded, and peptides were excluded from selection for 45 secs if fragmented n=2 times.

Crude peptide analysis was performed on a Q Exactive Plus hybrid quadrupole-orbitrap mass spectrometer coupled to an Agilent 1260 LC system to correct for variation in peptide loading across TMT channels using 2.5 kV no sheath or auxiliary gas flow; heated capillary temperature, 250°C. Approximately 30 ng of the supernatant from pTyr IP was loaded onto an in-house packed precolumn (100 um ID x 10 cm) packed with 10 μm C18 beads (YMC gel, ODS-A, AA12S11) connected in series to an analytical column (as previously described) and analyzed with a 75 min LC gradient [0-30% B from 0-40 mins, 30-60% B from 40-50 mins, 60-100% B from 50-55 mins, and 100-0% B from 60-65 mins]. MS1 scans were performed with m/z range: 350-2000; resolution: 70,000; AGC target: 3e^6^; max IT: 50 ms. The top 10 abundant ions were isolated (isolation width 0.4 m/z) and fragmented (nCE = 33%) with 70,000 resolution, max IT 150 ms, AGC target 1e^5^. Unassigned, +1, and >+7 charge states were excluded, and dynamic exclusion was set to 30 secs.

### MHC MS search space, filtering, and analysis

All mass spectra were analyzed with Proteome Discoverer (PD, version 2.5) and searched using Mascot (version 2.4) against the human SwissProt database (2020_06). No enzyme was used, precursor mass tolerance: 10 ppm, fragment mass tolerance: 20 mmu. Variable modifications were set to include oxidized methionine, static modifications included N-terminal and lysine TMT.

Heavy leucine-containing peptides were searched for separately with heavy leucine (+7), as a dynamic modification against a custom database of the synthetic peptide standards. All analyses were filtered with the following criteria: search engine rank =1, isolation interference ≤ 30%, ion score ≥ 15 and percolator q-value ≤ 0.05. Master protein descriptions were used to assign source proteins to ambiguous peptides for downstream analyses. Reporter ion intensities of PSMs assigned to the same peptide sequence were summed, and reporter ion intensities were corrected using hipMHC intensity values (CDK4/6i analysis only) as previously described.^7^ Only peptides with a length between 8 and 15 amino acids were considered for downstream analyses.

To evaluate differences between conditions, the log_2_ transformed ratio of arithmetic mean intensity for drug- and DMSO-treated samples (n=3) was calculated. To determine if peptides were significantly increasing/decreasing, an unpaired, 2-sided t-test was performed with p ≤ 0.05 set as the threshold for significance. PCA analyses were performed using Matlab R2019b.

### pTyr MS search space, filtering, and analysis

All mass spectra were analyzed with PD 2.5 and searched using Mascot 2.4 against the human SwissProt database (version 2020_06). For pTyr analyses, Spectra were searched using the following parameters: enzyme: trypsin, maximum missed cleavages: 2, precursor mass tolerance: 10 ppm, fragment mass tolerance: 20 mmu. Static modifications included TMT-10-labeled lysine and N-terminal residues, as well as cysteine carbamidomethylation. Dynamic modifications included methionine oxidation, and tyrosine, serine, and threonine phosphorylation.

Phosphorylation sites were localized with ptmRS module^15^ with 216.04 added as a diagnostic mass for pTyr the immonium ion.^16^ Peptides were filtered with the following criteria: search engine rank =1, isolation interference ≤ 35%, ion score ≥ 17, and ≥1 tyrosine phosphorylated residue. Peptides were filtered with the following criteria: search engine rank =1, isolation interference ≤ 35%, ion score ≥ 17, and ≥1 tyrosine phosphorylated residue. PSMs with >95% localization probability for all phosphorylation sites were classified as unambiguous and used for downstream analyses.

Crude peptide mixture was searched with the following parameters: enzyme: trypsin, maximum missed cleavages: 2, precursor mass tolerance: 10 ppm, fragment mass tolerance: 20 mmu. Static modifications included TMT-10-labeled lysine and N-terminal residues, as well as cysteine carbamidomethylation. Dynamic modifications included methionine oxidation. Peptides were filtered with the following criteria: search engine rank =1, ion score ≥ 20. Phosphotyrosine peptide reporter ion areas were corrected for variations in sample loading within each analysis using the median of peptide ratios in the crude peptide analysis for each channel relative to channel. Next, reporter ion intensities were summed across matching PSMs. Hierarchical clustering and PCA analyses were performed using Matlab R2019b.

### Peptide MHC binding affinity

Binding affinity of pMHCs was estimated using NetMHCpan-4.0 against the allelic profile of SKMEL5 cells.^17,18^ Only 9-mers were evaluated, and the minimum predicted affinity (nM) of each peptide was used to assign peptides to their best predicted allele. The threshold for binding was set at 500 nM.

### Enrichment analyses

For pMHC pathway enrichment analyses, gene names from peptide source proteins were extracted and rank ordered according to the average log_2_ fold change over DMSO treated cells. In cases where more than one peptide mapped to the same source protein, the maximum/minimum was chosen, depending on the directionality of enrichment analysis. We utilized gene set enrichment analysis (GSEA) 4.0.3 pre-ranked tool against the Molecular Signatures Database hallmarks gene sets with 1000 permutations, weighted enrichment statistic (p=1), and a minimum gene size of 15 for pMHC analyses.^19–21^ Results were filtered for FDR q-value ≤ 0.25, and nominal p-value ≤ 0.05.

### Experimental Design and Statistical Rationale

HipMHCs were titrated into 6 samples at 3 concentrations (n=2) to generate a 3-point calibration curve while minimizing protein input requirements. To compare 2 experimental conditions in the pMHC analyses (DMSO versus palbociclib treatment) and 3 experimental conditions (0s, 30s, 2m EGF stimulation) in pTyr analyses, n=3 biological replicates were selected for each condition to allow for calculating statistical significance.

## RESULTS

### Characterizing the quantitative accuracy of “boosted” pMHC analysis using synthetic, heavy isotope-labeled pMHCs

To interrogate the impact of including a carrier proteome on pMHC identification and quantitation, we prepared a set of 6 cell line-derived replicate samples comprised of 1×10^6^ cells per channel for the analysis without a protein carrier (“no-boost”), and a parallel experiment using 50% fewer cells per sample (5×10^5^ cells) for the “MHC-boost” analysis (**Fig. 1A, supplemental Fig. S1A**). As a protein carrier, we utilized 2 samples (2 channels) of 2.5×10^6^ cells stimulated with 10 ng/mL interferon-gamma (IFN-γ) for 72h. IFN-γ stimulation increases pMHCs levels ∼2-fold (**supplemental Fig. S1B-S1C**), resulting in a ∼10-fold boost per protein carrier sample and a combined signal-to-boost of ∼20-fold, in line with recent published guidelines for SCP experiments.^10^

**Fig. 1.**
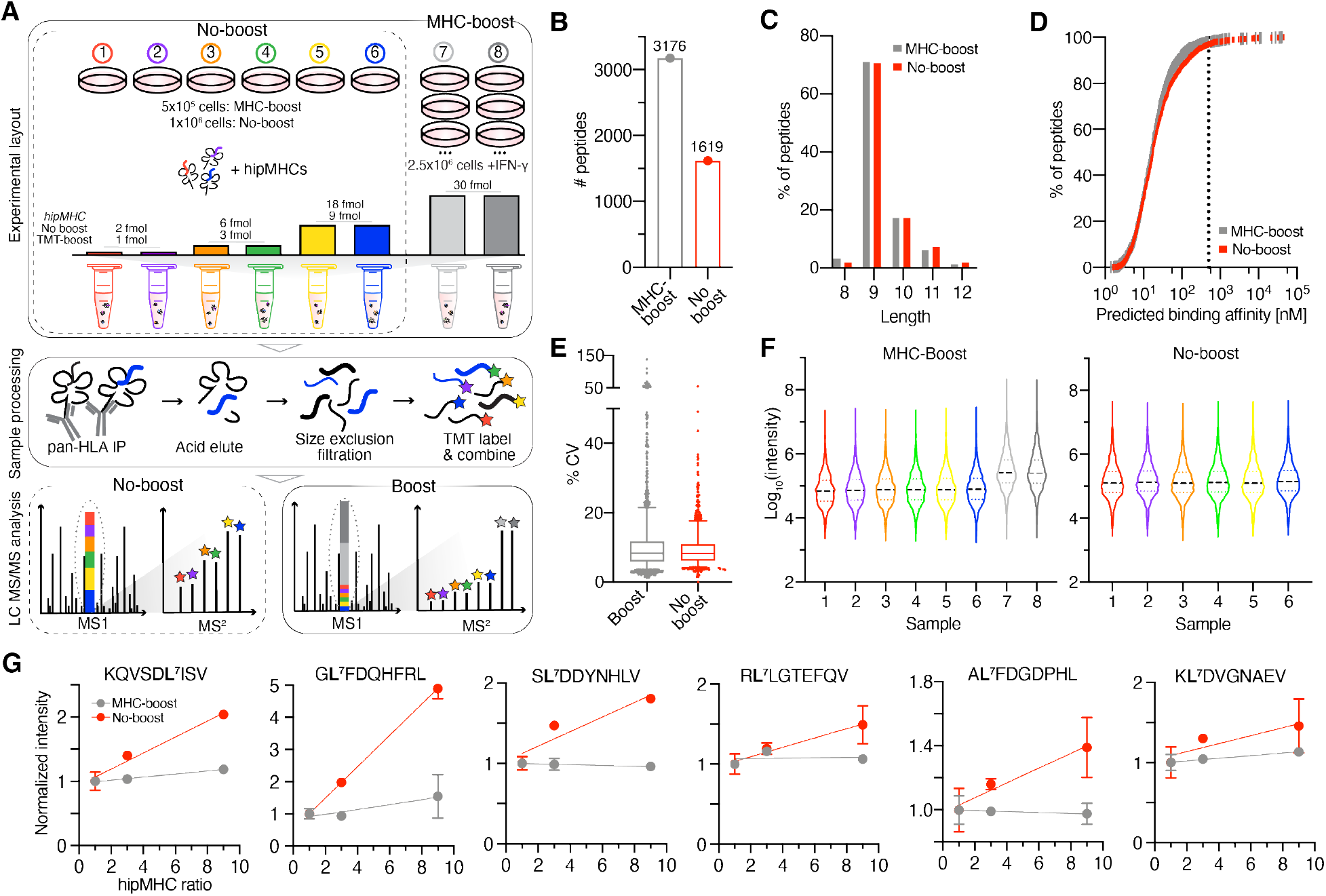
Estimating ion suppression using hipMHCs in immunopeptidomic analyses. A, Experimental setup of hipMHC quantitative immunopeptidomic analyses +/-protein carrier. B, Number of unique pMHCs identified in a single analysis. C, Length distribution of pMHCs. D, Predicted binding affinity of 9-mers. 97.9% and 97.0% of 9-mers in the pMHC-boost and no boost analyses, respectively, were predicted to have a binding affinity ≤ 500 nM (dotted line). E, Coefficients of variation of pMHC-boost and no-boost analyses. Boxes outline the interquartile range, and whiskers the 5 and 95th percentiles. pMHC-boost median CV= 8.23%, no-boost = 8.30%. 95% PSMs have CV <17.7% (no-boost) and 21.5% (pMHC-boost) F, Violin plots of reporter ion intensities for pMHC-boost (left) and no-boost (right) analyses. Median: black dashed line, quartiles: colored dotted line. G, Reporter ion intensities of hipMHC peptides normalized to the mean signal of the 1 fmol (pMHC-boost) or 2 fmol (no-boost) samples (y-axis), where the hipMHC ratio represents the amount of hipMHC added over the lowest concentration (x-axis). Solid line = linear fit, error bars show +/-standard deviation.

To measure ion suppression, we utilized a panel of six synthetic, heavy-isotope labeled pMHCs (hipMHCs), which were titrated into cell lysates prior to pMHC isolation to generate an internal standard curve against a consistent background immunopeptidome, as previously described^7^. HipMHCs were added at a ratio of 1:1:3:3:9:9 across the 6 samples, with concentrations of 1, 3, and 9 fmol in the boost analysis, and proportionally, 2, 6, and 18 fmol in the no-boost analysis. The protein carrier samples contained 30 fmol of each hipMHC, 10-fold more than the median concentration used across non-protein carrier samples (**supplemental Fig. S1A**). After addition of hipMHCs, class I pMHC complexes were isolated from each sample by immunoprecipitation, acid elution, and size exclusion filtration. Peptides for each sample were subsequently labeled with TMT, combined, and analyzed by LC-MS/MS.

As expected, including a protein carrier resulted in a large increase in the number of unique pMHC IDs using 50% less cellular input material for each channel: from a single injection using just 25% of the labeled mixture, 3176 unique pMHCs were identified in the pMHC-boost sample, whereas 1619 were identified in the no-boost analysis (**Fig. 1B**). The peptides identified in both experiments followed expected length distributions (**Fig. 1C)**, with 97.0% and 97.9% of 9-mers predicted to be allelic binders in no-boost and pMHC-boost analyses, respectively (**Fig. 1D**). While both analyses had equivalent median coefficients of variation (CV) across replicates (**Fig. 1E**), peptide spectrum matches (PSMs) in the boost analysis had a wider distribution of CV values. Together, these data suggest that a 20x protein carrier improves the number of unique IDs while not altering peptide properties of the resultant data set but may result in slightly higher quantitative variation. Of note, the proportion of missing values between the protein carrier and non-carrier samples in the pMHC boost analysis were comparable (4% of PSMs in no-boost, 8% in pMHC-boost), suggestive of sufficient ion sampling for a majority of peptides (**supplemental Fig. S1D)**.

We next examined the intensity distributions across PSMs and found that the protein carrier samples had 3.5-4-fold higher intensity than the other samples in the boost analysis (**Fig. 1F**). Our expected intensity ratios were ∼10:1 (5-fold increase in sample in the protein carrier channels, coupled to a 2-fold increase in MHC expression due to IFN-γ), and thus the observed peptide ratios demonstrate approximately a 60% reduction in signal intensity, suggestive of ion suppression. Ratios of the titrated hipMHCs were subsequently analyzed, and substantial ion suppression was observed in both analyses (**Fig. 1G**). For example, in the no-boost analysis the “GLFDQHFRL” peptide had a 1.8-fold reduction in dynamic range, while the “KLDVGNAEV” peptide had a 6.2-fold reduction, with the other hipMHC peptides falling between these two extremes. While the hipMHC intensity ratios did not match expected values in the no-boost analysis, reporter ion intensities did increase with increasing concentration of hipMHC. By comparison, the quantitative accuracy in the MHC-boost analysis was severely negatively affected by the presence of the protein carriers, as there was minimal difference in the reporter ion intensities for the hipMHC standards across all samples, with “GLFDQHFRL,” being the only exception (6.7-fold reduction in observed vs. expected dynamic range). Taken together, these data demonstrate that while ion suppression exists in non-boost and boost experiments alike, the presence of a protein carrier increased ion suppression to the extent that pMHCs up to 9-fold higher in concentration could not be differentiated via isobaric intensities. It is worth noting hipMHCs were added at relatively high concentrations, representing a range of ∼1000-10,000 pMHCs/cell. Quantitative accuracy of endogenous pMHCs at lower presentation levels may be further negatively impacted by the presence of a protein carrier.

### Protein carrier channel skews biological interpretation of palbociclib-induced pMHC repertoire alterations

To further assess the accuracy of quantifying endogenous pMHCs in the presence of a protein carrier in a biological context, we evaluated whether a carrier proteome would affect data interpretation of melanoma cells treated with the CDK4/6 inhibitor, palbociclib, which increases pMHC presentation and induces palbociclib-specific repertoire changes, as previously reported.^7^ Cells were treated with 10 µM palbociclib or DMSO as a vehicle control for 72h in triplicate, and analyzed alone or with an IFN-γ stimulated protein carrier channel for a combined 20-fold signal-to-boost ratio, using a similar set-up to the previous experiment. (**Fig. 2A, supplemental Fig. S2A**).

**Fig. 2.**
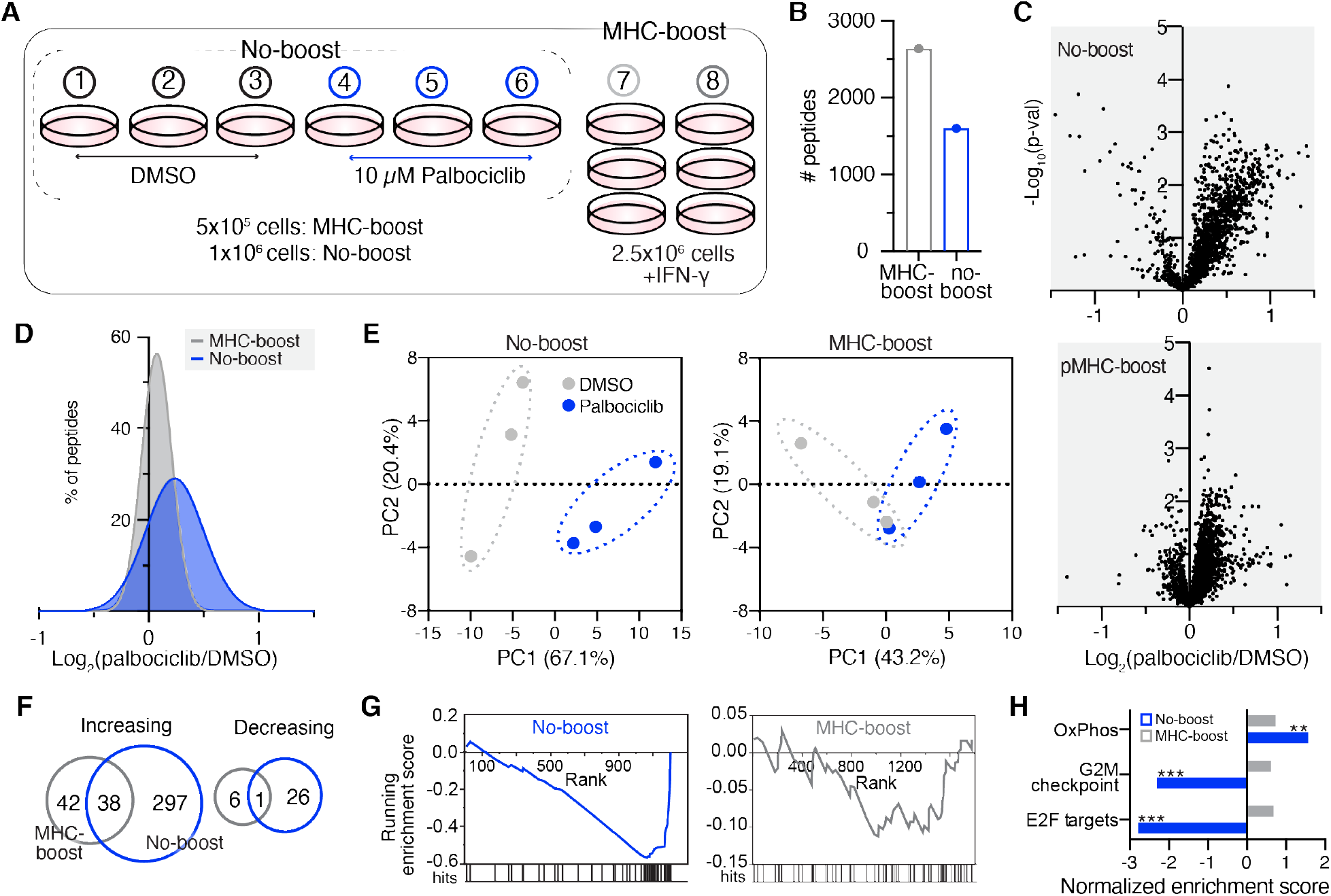
Palbociclib-induced pMHC repertoire alterations are masked in the presence of a protein carrier. A, Experimental setup of pMHC analyses +/-protein carrier with 72h DMSO or 10 µM palbociclib treatment. B, Number of unique peptides identified in pMHC-boost (2637) and no-boost (1602) analyses. C, Volcano plot displaying the log_2_(palbociclib/DMSO) of pMHCs (x-axis), where the fold change is calculated from the mean intensity of n = 3 biological replicates per condition, versus significance (y-axis, mean adjusted p-value, unpaired two-sided t test). D, Histogram distribution of unique pMHC fold change in expression. E, Samples plotted by principal component 1 (PC1) and PC2 score for no-boost (left) and pMHC-boost (right) analysis, colored by treatment condition. Percentages are % variance explained by the plotted PC. F, Venn diagram of peptides significantly increasing (upper) and decreasing (lower) with palbociclib treatment in the no-boost (blue) and pMHC-boost (grey) analyses. G, pMHC enrichment plots for E2F targets for the no-boost (grey, p=0.13, q=0.88) and pMHC-boost (blue, p<0.001, q<0.001) analyses. Hits mark pMHCs of source proteins mapping to E2F targets. H, Normalized enrichment scores from enrichment analyses of pMHC-boost (grey) and no-boost (blue) datasets. Positive/negative scores represent directionality of pathway enrichment. Significant enrichment is noted by **p<0.01, ***p<0.001, with FDR-q values < 0.25.

Similar to the hipMHC experiment, ‘boosting’ with a protein carrier yielded a greater number of unique peptides identified (2637 in the “MHC-boost” analysis vs. 1602 in the “no-boost” analysis) (**Fig. 2B)**, with similar length distributions (**supplemental Fig. S2B**). The no-boost experiment recapitulated our previously reported results^7^, where a majority of peptides showing an slight increase in presentation levels following palbociclib treatment (median fold change 1.17x), while peptides in the MHC-boost experiment showed a narrower distribution of changes, centered around a median fold change of just 1.05x (**Fig. 2C-2D)**. In line with this finding, principal component analysis (PCA) showed superior separation of DMSO and palbociclib-treated samples in the no-boost versus the MHC-boost analysis (**Fig. 2E)**.

To interrogate the data further, we considered the 1092 unique peptides quantified in both analyses (**supplemental Fig. S2C**). Of these peptides, fewer peptides were significantly increasing or decreasing in presentation in the MHC-boost analysis compared to the non-boost analysis (**Fig. 2F**), masking biological interpretation of the data. For example, 334 common peptides significantly increased in presentation in the non-boost analysis, while only 80 common peptides in the boost analysis significantly increased.

Interestingly, 42 of the 80 peptides were significantly increased in only the pMHC-boost but not the no-boost analysis. Upon closer inspection, we found 76% of peptides also showed in increase in presentation in the no-boost analysis but did not achieve statistical significance. Ion suppression can reduce variation in reporter-ion intensities, which we observed as reduced median coefficients of variation in the MHC-boost analysis compared to the no-boost analysis (**supplemental Fig. 2D**). This may artificially increase the likelihood of statistical significance among replicate samples, offering a possible explanation for this finding.

We next evaluated whether the altered quantitation in the boost analysis would change the previously described key findings of this experiment, namely that MHC peptides derived from proteins in pathways known to be perturbed by CDK4/6 inhibition show significant positive enrichment (oxidative phosphorylation, OxPhos) and negative enrichment (G2M checkpoints and E2F targets).^7^ To this end, we performed an enrichment analysis using the MSigDB Hallmarks gene set database by rank ordering the gene names for pMHC source proteins in decreasing order of fold-change.^19–21^ In the no-boost analysis, 10 µM palbociclib treatment showed significant enrichment in OxPhos, G2M checkpoints, and E2F targets, mirroring previously reported findings (**Fig. 2G-2H**). In contrast, no pathways, including the three highlighted in the no-boost analysis, showed significant enrichment using the pMHC-boost dataset data. A comparison of E2F target peptides between the analyses illustrates this finding—most peptides with decreased expression in the no-boost analysis showed little change in expression in the presence of a protein carrier. (**supplemental Fig. S2E**). These data reaffirm that while utilizing a protein carrier channel can increase the number of peptides identified and quantified across samples using lower cellular input, ion suppression due to the presence of a protein carrier can alter quantitative dynamics to the extent that known biological findings are masked, hiding relevant insight.

### Effects of PV-stimulated protein carrier on quantitative phosphotyrosine analyses

Since the effect of boosting appeared to adversely affect quantitative accuracy in the immunopeptidomics experiments, we sought to evaluate whether utilizing a protein carrier would also impact quantitative accuracy in pTyr analyses. To provide a set of samples with altered signaling of a biologically relevant network for quantification, we utilized A549 cells stimulated with 5 nM EGF for 0 seconds, 30 seconds, or 2 minutes (0s, 30s, 2m) to drive a dynamic response in tyrosine phosphorylation levels among a subset of epidermal growth factor receptor (EGFR)-related pTyr sites, as previously described.^12,14,22^ Three biological replicates of 100 µg input material for each time point were utilized in the “no-boost” analysis, whereas the “PV-boost” analysis contained the same replicate samples along with 1 mg of protein carrier, A549 cells stimulated with PV to halt tyrosine phosphatase activity, thereby driving elevated pTyr signal (**Fig. 3A**). Peptide amounts were selected to match the upper and lower limits of sample input utilized by a previously reported pTyr boosting study (Chua et al.), however we utilized a lower concentration of pervanadate (30 µM versus 500 µM).^5^ Following tryptic digestion & standard sample processing, samples were labeled with TMT-10plex, and tyrosine phosphorylated peptides were subsequently purified using two-step enrichment followed by LC-MS/MS analysis (**Fig. 3A-3B**).^14,22^

**Fig. 3.**
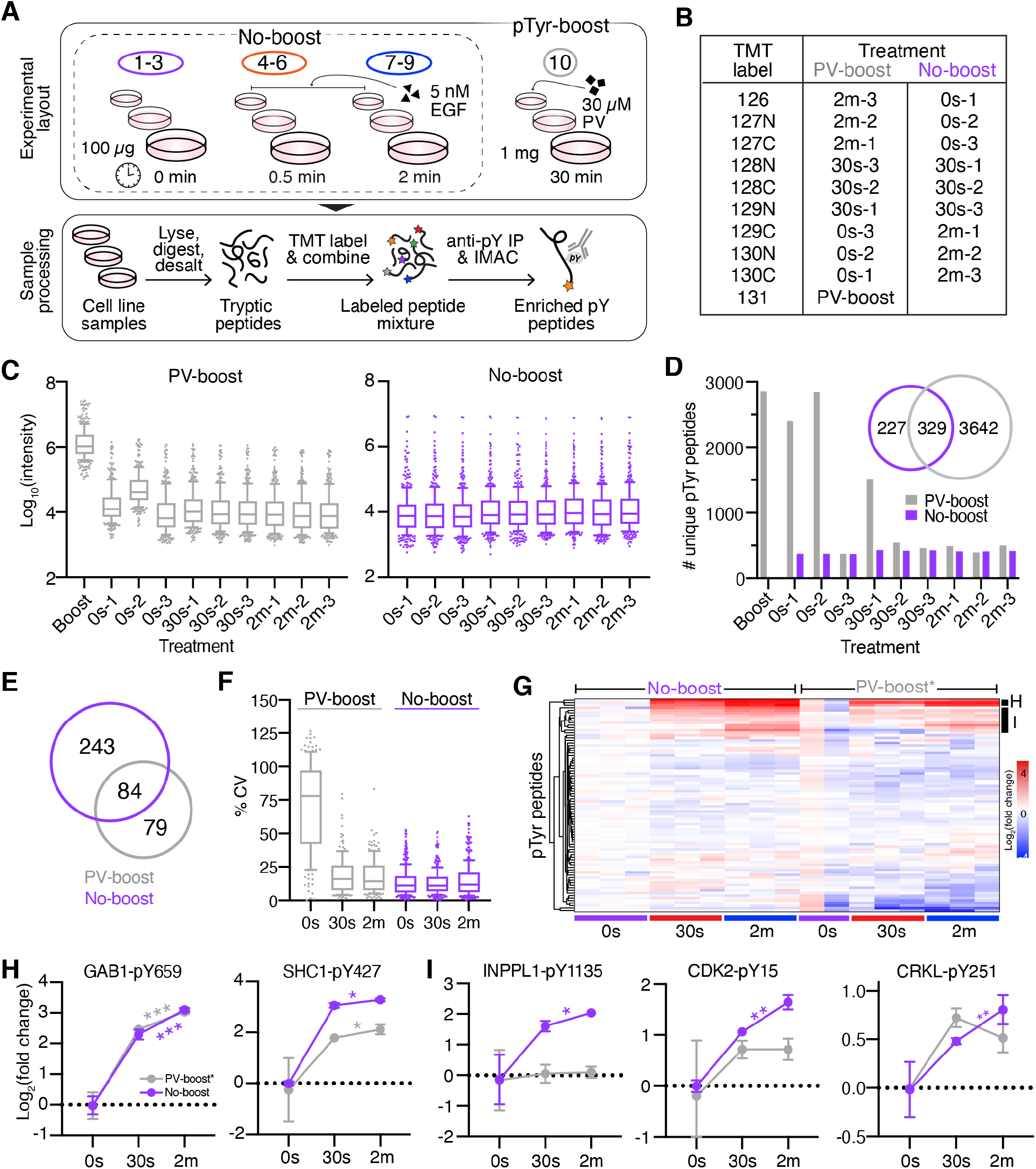
Analysis of EGF-induced signaling dynamics in the presence of a PV-treated protein carrier. A, Schematic of PV-boost vs. no-boost experimental layout and pTyr peptide enrichment. B, Isobaric labeling scheme and treatment conditions. C, Reporter ion intensities for PV-boost (left) and no-boost (right) analyses. Boxes outline the interquartile range, and whiskers the 10 and 90th percentiles. D, Number of unique pTyr peptides quantified in each sample. Venn diagram shows overlap in total pTyr peptides between analyses. E, Venn diagram of the number of unique pTyr peptides quantified across all samples in each analysis (no missing values). F, Coefficients of variation of PV-boost (grey) and no-boost (purple) analyses. Boxes outline the interquartile range, and whiskers the 10 and 90th percentiles. PV-boost median CV: 14-78%, no-boost: 11-12%. G, Hierarchical clustering (Euclidean) of Log_2_(fold change) values of pTyr sites identified in both analyses. PV-boost* values represent the original values from the PV-boost analysis re-normalized to the mean 0s intensity with the 0s-2 excluded. H-I, Log_2_(fold change) values of peptides in clusters H-I (Figure 3G). Significance values: 2-tailed t-test of 30s vs. 2m timepoint. *p<0.05, **p < 0.01, ***p<0.001. Error bars represent +/-standard deviation.

Even though we treated cells with a lower concentration of PV relative to Chua et al., the PV-treated protein carrier sample still had substantially higher reporter ion intensities (∼100-fold) compared to the EGF-stimulated samples, well outside the suggested protein carrier-to-signal range recommended for SCP boost experiments (**Fig. 3C**). Indeed, the high signal level from the TMT-131 labeled PV-boost protein carrier channel resulted in isotopic interference in the second replicate of the zero-second channel (0s-2) labeled with the 130N TMT tag. By comparison, the no-boost analysis showed similar intensity distributions across samples.

As anticipated, the PV-boost analysis identified a considerably higher number of unique pTyr peptides compared to the no-boost analysis (3971 vs. 556) (**Fig. 3D**). However, a majority of identified peptides in the PV-boost analysis were only quantified in the protein carrier channel or adjacent channels (isotopic interference), resulting in a large number of PSMs with missing values (up to 94%). By comparison, the no-boost analysis had far fewer peptides with missing values (up to 17%). Consequently, despite the greater number of overall pTyr-peptide identifications, the PV-boost analysis contained just 163 pTyr peptides quantifiable across all samples versus 327 in the no-boost analysis, reducing overall data quantity by 2-fold (**Fig. 3E**). The number of EGFR signaling related peptides was similarly reduced with 40 vs. 20 pTyr peptides mapping to proteins in KEGG ErbB signaling pathway in the no-boost and PV-boost analyses, respectively.^23^

To assess whether the PV-treated protein carrier channel also influenced the accuracy of the quantitative temporal signaling data, we compared the coefficients of variation between analyses (**Fig. 3F**). The 30s and 2m timepoints showed slightly higher variability in the PV-boost analysis versus the no-boost analysis, where for example the median 30s CV was 11% in the no-boost analysis compared to 16% in the PV-boost analysis. The 0s timepoint in the PV-boost analysis exhibited high CVs as a result of the isotopic interference (median 78%), greatly altering quantitative accuracy.

To compare the quantitative dynamics between analysis, a hierarchical clustering analysis of the 84 peptides quantified in both analyses was performed (**supplemental Fig. S3A**). The 0s-2 sample with isotopic interference greatly skewed quantitation by increasing the mean 0s signal, thus most of the peptides in the PV-boost analysis appear to have decreased phosphorylation in response to EGF, as compared to the no-boost analysis where the same sites show constant phosphorylation levels (**supplemental Fig. S3B**). Moreover, the increase in the mean 0s quantitation in the PV-boost analysis resulted in substantial ratio compression among peptides modulated by EGF stimulation (**supplemental Fig. S3C-S3D**).

To better assess the effects of the PV-boost protein carrier on quantitative accuracy, we removed the 0s-2 data labeled with 130N, after which the quantitative dynamics more closely mirrored those of the no-boost condition (**Fig. 3G**). Several of the EGF-modulated peptides showed a large increase in phosphorylation after stimulation, and while a few peptides showed correlated dynamics like GAB1-pY659, others still showed dynamic range suppression (**Fig. 3H**). For example, we measured an 11-fold increase in pTyr for SHC1-pY427 following 2 minutes of EGF stimulation in the no-boost analysis, which was reduced to a 4-fold change when analyzed with a protein carrier. While the same trend of increased phosphorylation with EGF stimulation was preserved between the analyses for the SHC1 peptide, subtler pTyr changes may be masked by the effects of ion suppression. This was seen in the INPPL1-pY1135, CDK2-pY15 and CRKL-pY251 peptides, which have significantly different pTyr levels between the 30s and 2m timepoints in the no-boost analysis but are not significantly different in the PV-boost dataset (**Fig. 3I**). Indeed, 33 peptides have significantly different pTyr levels (p<=0.05) between the 30s and 2m condition in the no-boost analysis versus just 20 in the PV-boost analysis (**supplemental Fig. S4A)**, indicative of increased dynamic range suppression. PCA reinforces this finding, as the 30s and 2m sample cluster closer together (regardless of inclusion/exclusion of the 0s-2 sample), whereas the no-boost samples cluster with superior separation (**supplemental Fig. S4B**).

### Reduction in protein carrier improves quantitative accuracy but still increases missing values

To determine whether we could improve quantitative accuracy and overall data quality by using a smaller amount of a more targeted boost channel, we performed two additional experiments using 100 µg in triplicate of the 5 nM EGF stimulated samples at 0, 30s, and 2m time points used in the PV-boost/no-boost experiments along with a 900 µg protein carrier consisting of equal parts of each sample for a boost-to-signal ratio of approximately 9-fold (“EGF-boost”) (**Fig. 4A-4B**). Unlike the PV-boost, which inhibits tyrosine phosphatases and results in a general increase in most, if not all, phosphorylated tyrosines, we hypothesized that using EGF-stimulated samples as a boost would lead to more targeted detection of the EGFR signaling network. Additionally, to assess whether increased ion numbers might yield improved quantitative accuracy and fewer missing values, the EGF-boost analyses were analyzed under two conditions: at an AGC target of 1e^5^ and maximum IT of 250 ms, as performed in the PV-boost/no-boost analyses, and with an increased AGC target of 5e^5^ and maximum IT of 500 ms.

**Fig. 4.**
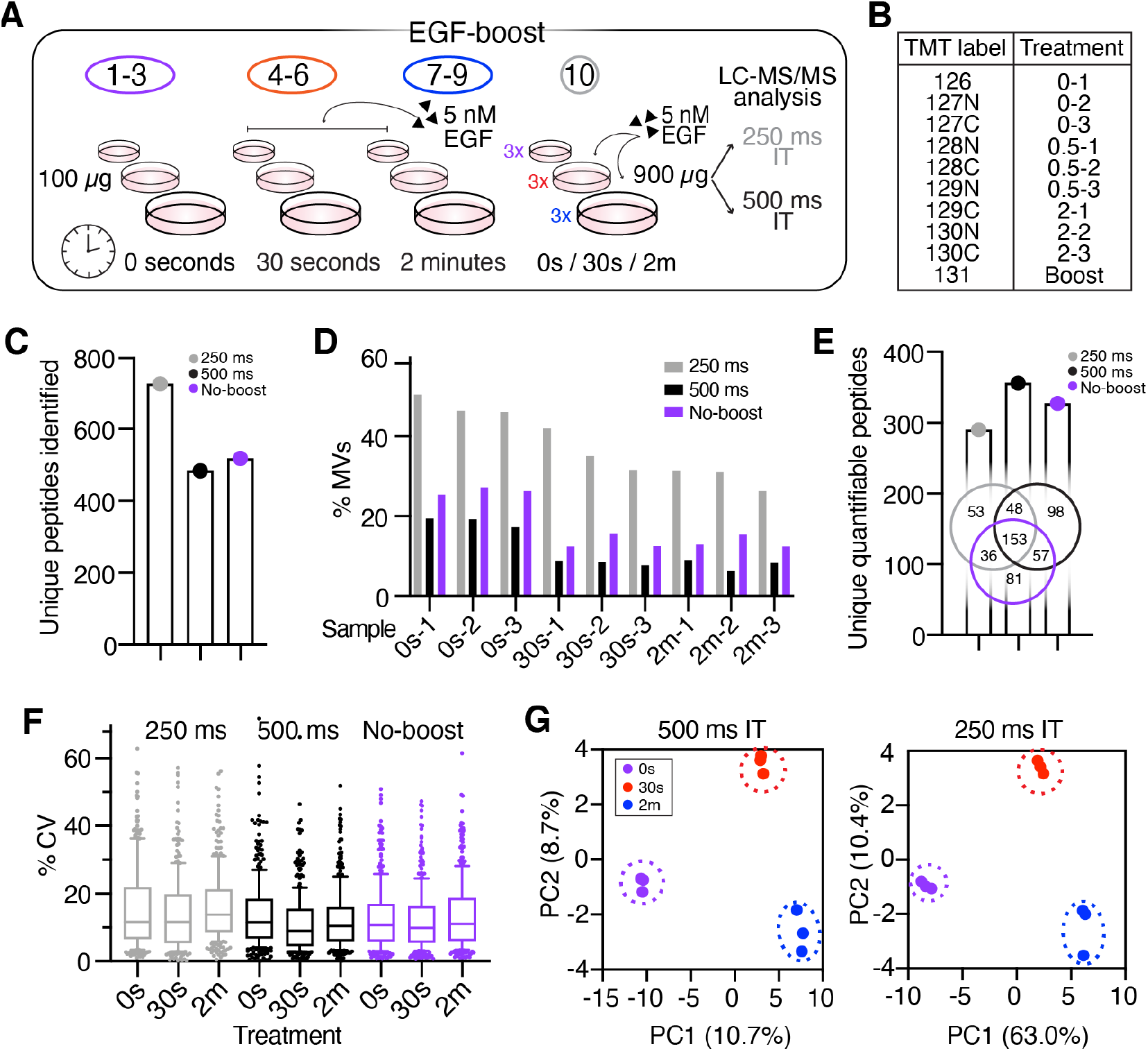
pTyr-boost with 9x EGF-boost protein carrier improves quantitative accuracy but still yields large number of missing values. A, Experimental layout of EGF-boost experiment with 9x protein carrier. B, Isobaric labeling scheme and treatment conditions. C, Total number of unique pTyr peptides identified in each analysis. D, Proportion of PSMs with missing values for each sample. E, Total number of unique pTyr peptides quantified in each analysis and Venn-diagram of peptides commonly identified between analyses (no MV’s). F, Coefficients of variation between replicates in EGF-boost and no-boost analyses. Boxes outline the interquartile range, and whiskers the 10 and 90th percentiles. EGF-boost 250 ms median CV: 12-14%, 500 ms IT: 9-12%, no-boost: 10-11%. G, Samples plotted by principal component 1 (PC1) and PC2 score, colored by EGF stimulation condition for 500 ms EGF boost and no-boost analyses. Percentages describe the variance explained by the plotted PC.

As expected, an increased number of unique pTyr peptides were identified in the 250 ms analysis compared to the 500 ms analysis (**Fig. 4C, supplemental Fig. S5A**). However, the proportion of PSMs with MVs was similarly increased in the 250 ms analysis (250 ms: 26-50% MVs, 500 ms: 6-19% MVs), resulting in fewer unique pTyr peptides quantifiable across all samples in the 250 ms analysis (290) versus the 500 ms analysis (356) (**Fig. 4D-E**). In comparison to the 327 pTyr peptides identified and quantified in the no-boost analysis previously described, the 500 ms IT “EGF boost” offers a slight increase in data quantity. In both EGF-boost analyses, we identified 45 EGFR-related peptides quantified across all samples, representing a slight improvement over the no-boost data (40) and more than double the EGF-related peptides identified in the PV-boost data. This finding is in support of our hypothesis that an EGF-stimulated protein carrier would enhance data quantity of EGF signaling-related peptides.

We next compared intensity distributions for each of the 9 EGF stimulated samples to evaluate data quality and found them to be similar in both EGF-boost analyses with no obvious isotopic interference from the protein carrier, which had an increased intensity distribution near the expected 9-fold ratio (**Figure S5B**). We verified this by comparing the CVs between replicates and found that the 500 ms IT and no-boost analyses had comparable median CVs (10.4% and 10.6%, respectively), whereas the 250 ms IT analysis had a lightly higher median CV (12.4%) (**Figure 4F)**. Nevertheless, a PCA analysis showed clear separation of samples by treatment condition in both EGF-boost analyses, a substantial improvement over the PV-boost analysis (**Figure 4G**).

To further assess quantitative accuracy, the 153 peptides commonly identified and quantified across the three analyses were hierarchically clustered (**Figure 5A**), displaying similar patterns of phosphorylation with EGF-responsive peptides clustering together (**Figure 5A and S5C)**. While some of the peptide had significantly correlated phosphorylation dynamics between the no-boost and EGF-boosted analyses (**Figure 5B**), others showed significant correlation only in the no-boost and 500 ms IT condition (**Figure 5C**). Despite this finding, even in the 500 ms dataset, fewer peptides showed a significant change in phosphorylation from the 0s control at the 30s and 2m timepoints compared to the no-boost analysis (**Figure S5D**). Of the peptides that were not significant in the 500 ms analysis, there was evidence of dynamic range suppression and altered quantitative dynamics (**Figure 5D**), highlighting that not all peptides had comparable quantitation between the boost and non-boost analyses.

Together, these data demonstrate that a smaller and more targeted carrier-to-signal ratio may improve quantitative accuracy compared to a larger protein carrier. However, the smaller protein carrier offers only a slight benefit in data quantity and still demonstrates reduced quantitative accuracy compared to the no-boost control, even when using instrument parameters designed to improve accuracy.

**Fig. 5.**
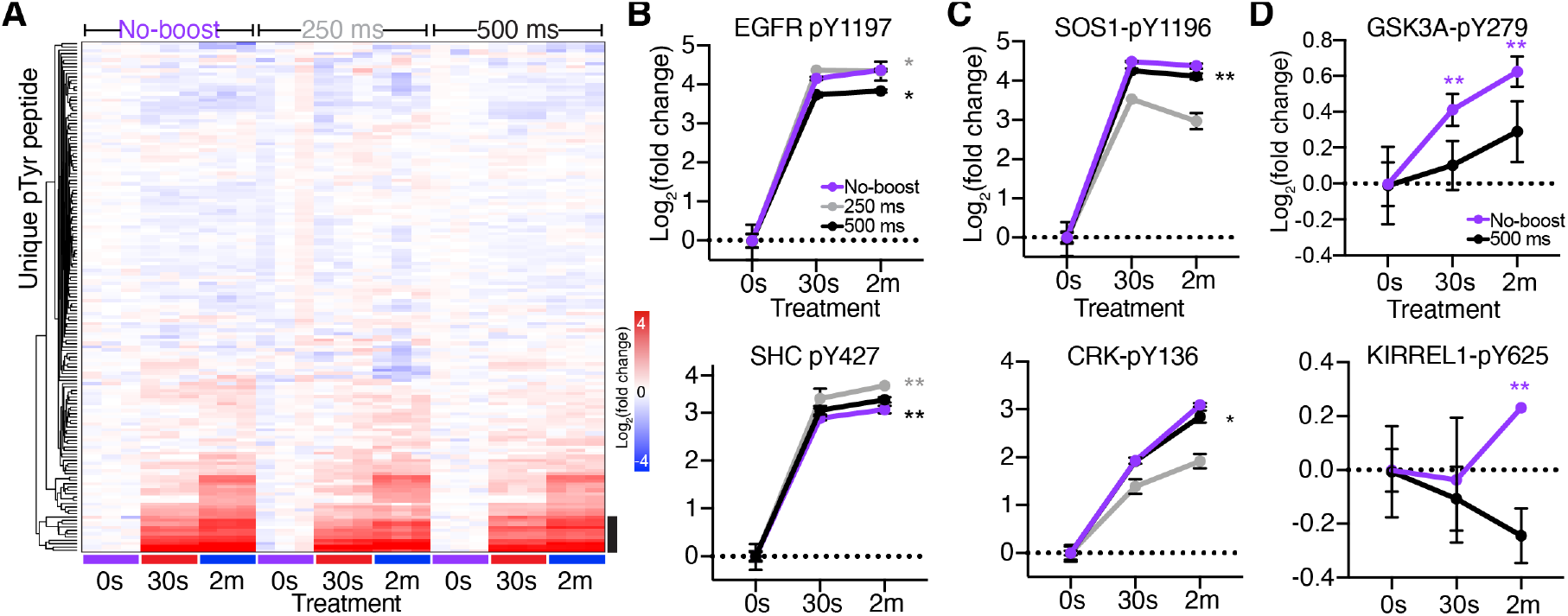
9x EGF-stimulated protein carrier has higher quantitative accuracy, but EGF-modulated sites still show altered pTyr dynamics and ratio compression. A, Hierarchical clustering of peptides identified in all three analyses, represented as log_2_(fold change) of each sample normalized to the mean reporter ion intensity of the 0s control per analysis. Black bar highlights EGF-modulated peptides highlighted in B-C, supplemental Fig. S5C. B-C, Log_2_(fold change) of pTyr signal of selected peptides from clusters B and C highlighted in 5A. Error bars represent +/-standard deviation (n=3). Pearson correlation significance (two-tailed): *p<0.05, **p<0.01. D, Log_2_(fold change) pTyr signal of peptides. Significance values: 2-tailed t-test of 0s vs. 30s/2m timepoint. **p<0.01, Error bars represent +/-standard deviation.

## DISCUSSION

The ability to reduce sample input and/or increase signal with a protein carrier is particularly appealing in immunopeptidomics and tyrosine phosphoproteomics, two applications that are often limited by larger sample requirements. However, our data indicate that inclusion of a protein carrier decreases quantitative accuracy in MS^2^-based quantitative analyses, even when using a signal-to-boost ratio within SCP guidelines (20x).^10^ Loss of quantitative accuracy associated with ‘boosting’ manifested as high ratio compression in pMHC analyses that masked dynamic alterations in pMHC expression levels and obscured known biological findings. Ratio compression was similarly observed in pTyr analyses, with the degree of ratio compression amplified with increasing signal in the protein carrier channel. To offset ratio compression and thus improve quantitative accuracy in ‘boosted’ sample analyses, triple-stage mass spectrometry (MS^3^) and/or high-field asymmetric waveform ion mobility spectrometry (FAIMS) have been shown to reduce ratio distortion, although both methods can come at a cost of sensitivity and data quantity.^24,25^ Additional experiments, similar in format to those described here, will be useful in determining whether MS^3^ and/or FAIMS can offer improved quantitative accuracy without compromising data quantity in this setting. To enable such comparisons, hipMHCs provide a useful tool to evaluate ion suppression in place of exogenously added peptide standards. It is worth noting that while MS^3^ may be applicable to improve quantitative accuracy for pMHC analysis, it is a relatively unattractive solution for tyrosine phosphoproteomics due to the cost in sensitivity, lower precision, and fewer peptide identifications compared to MS^2^.^26^

In pTyr analyses, utilizing a PV-treated protein carrier provided a strong increase in MS1 signal and greatly increased the number of pTyr peptides identified. Unfortunately, large proportions of missing values in this analysis decreased the overall data quantity compared to a parallel analysis performed without the protein carrier. While Chua et al. were able to replace missing values by interpolation, this strategy is not applicable for analyzing biological systems where the quantitative dynamics are unknown. In addition to missing values, the high signal level of the PV-boost protein carrier resulted in isotopic interference in adjacent channel(s) that negatively impacted quantification of these channels and their respective conditions. These channels could be removed in post-processing to improve quantitative accuracy, but this approach decreases the number of TMT tags available for sample multiplexing, diminishing the throughput and utility of this approach. Furthermore, dynamic range suppression was still observed even after excluding the sample with highest isotopic interference, suggesting that a boost-to-signal ratio of this magnitude may adversely affect the quantitative accuracy of the experiment regardless of isotopic leakage.

Decreasing the magnitude of the protein carrier in pTyr analyses and increasing the maximum IT and AGC target decreased MVs compared to the PV-boost analysis and increased the total number of identified and fully quantified peptides by 29 compared to the no-boost analysis. Despite this slight improvement in quantifiable phosphopeptides, some peptides still showed altered dynamics and ratio compression relative to the no-boost analysis, suggesting that use of a protein carrier in this experimental design is of little benefit. Further increasing the AGC target/IT may improve quantitative accuracy but will likely reduce the number of scans acquired and thus the number of identified peptides, though the balance between data quantity and data quality remains to be thoroughly explored. Alternatively, decreasing the number of multiplexed samples would increase ion sampling of non-carrier samples, but limited multiplexing would further reduce the utility of the assay.

These data illustrate that experiments leveraging protein carriers should rigorously evaluate the quantitative impact of the protein carrier (namely ion suppression, missing values, coefficients of variation, and isotope leakage) to avoid misinterpretation of biological data. Future studies exploring alternative instrument acquisition parameters and configurations, along with protein carrier magnitudes and signal stimulation strategies, will further illuminate whether protein carriers can be effectively used for quantitative studies in these applications, or whether improvements in sample preparation and instrument sensitivity may pave an alternative path forward in achieving high accuracy, high precision measurements without a signal boost.

## ACKNOWLEDGEMENTS

This research was supported in part by NIH grants R01EB027717, U54CA210180, and U01CA238720, as well as funding from MIT Center for Precision Cancer Medicine. L.E. Stopfer is supported by an NIH Training Grant in Environmental Toxicology (# T32-ES007020).

## CONTRIBUTIONS

LS and JCP conceived the project, performed experiments, analyzed and interpreted the data, and wrote the manuscript. FMW advised the project and assisted with manuscript writing and revisions.

