## Supplementary Data for "Quantitative consequences of protein carriers in immunopeptidomics and tyrosine phosphorylation MS^2^ analyses"

Supplemental Data

### SUPPLEMENTAL FIGURES

#### Figure S1

**A**

| TMT label | Cell # | | hipMHC (fmol) | |
|---|---|---|---|---|
| | No-boost | MHC-boost | No-boost | MHC-boost |
| 126 | $1e^6$ | $5e^5$ | 2 | 1 |
| 127N | $1e^6$ | X | 2 | X |
| 127C | X | $5e^5$ | X | 1 |
| 128N | X | $5e^5$ | X | 3 |
| 128C | $1e^6$ | X | 6 | X |
| 129N | $1e^6$ | $5e^5$ | 6 | 3 |
| 129C | X | $5e^5$ | X | 9 |
| 130N | X | $5e^5$ | X | 9 |
| 130C | $1e^6$ | $2.5e^6$ + IFN-γ | 18 | 30 |
| 131 | $1e^6$ | $2.5e^6$ + IFN-γ | 18 | 30 |

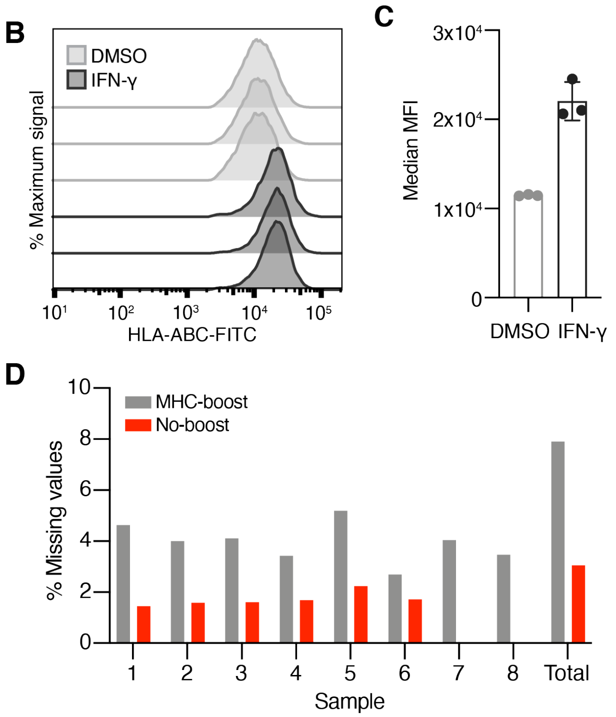


**Supplemental Fig. S1.**

*A,* Experimental setup of isobaric labels, cell number, and concentration of hipMHC added for each sample.

*B,* Flow cytometry analysis of surface HLA expression in SKMEL5 cells after 72h of DMSO or IFN-g treatment. Data are presented as % of maximum fluorescence intensity signal. N=3 biological replicates per treatment condition.

*C,* Median MFI of flow cytometry measurements in B. Error bars represent standard deviation from n=3 biological replicates.

*D,* Percentage of total PSMs with missing values in each sample.

**Figure S2**

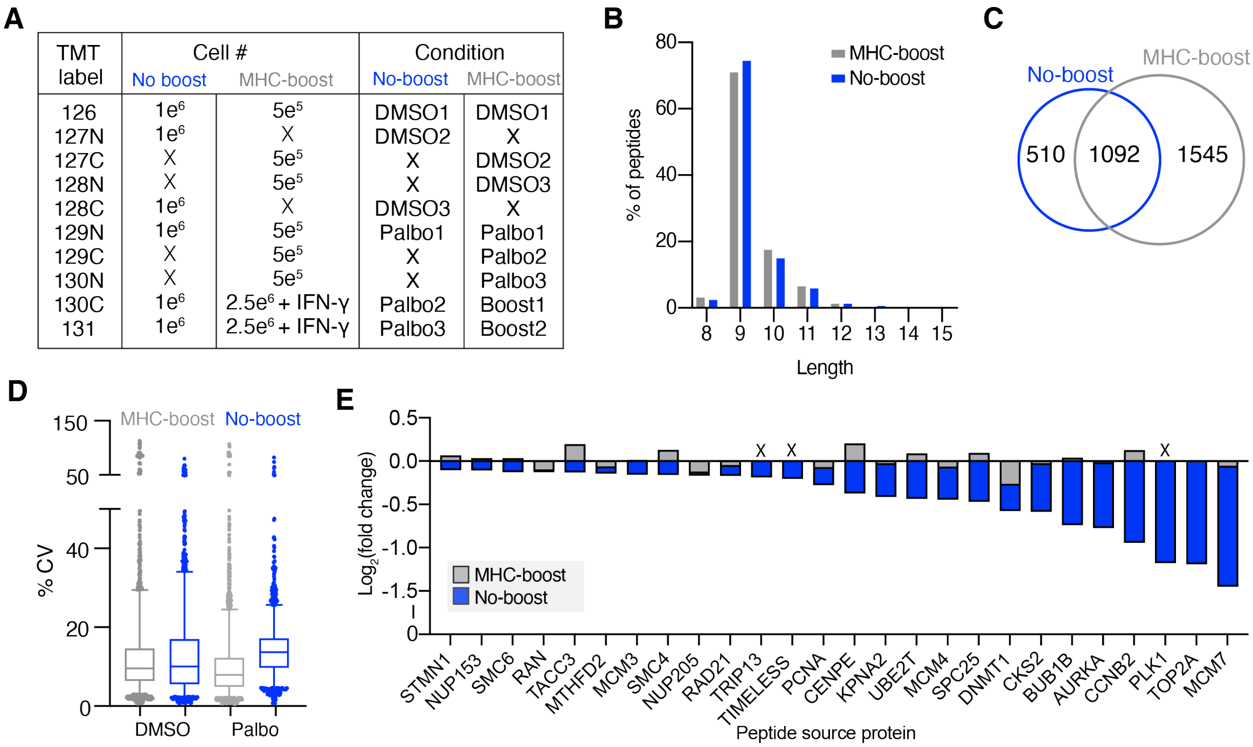


| TMT label | Cell # No boost | Cell # MHC-boost | Condition No-boost | Condition MHC-boost |
|---|---|---|---|---|
| 126 | $1e^6$ | $5e^5$ | DMSO1 | DMSO1 |
| 127N | $1e^6$ | X | DMSO2 | X |
| 127C | X | $5e^5$ | X | DMSO2 |
| 128N | X | $5e^5$ | X | DMSO3 |
| 128C | $1e^6$ | X | DMSO3 | X |
| 129N | $1e^6$ | $5e^5$ | Palbo1 | Palbo1 |
| 129C | X | $5e^5$ | X | Palbo2 |
| 130N | X | $5e^5$ | X | Palbo3 |
| 130C | $1e^6$ | $2.5e^6$ + IFN-γ | Palbo2 | Boost1 |
| 131 | $1e^6$ | $2.5e^6$ + IFN-γ | Palbo3 | Boost2 |

**Supplemental Fig. S2.**

*A,* Experimental setup of isobaric labels, cell number, and treatment conditions.

*B,* Length distribution of pMHCs.

*C,* Venn diagram of unique pMHC identified in the no-boost (blue) and pMHC-boost (grey) analysis.

*D,* Coefficients of variation of pMHC-boost and no-boost analyses. Boxes outline the interquartile range, and whiskers the 5 and 95th percentiles. pMHC-boost median CV: DMSO= 9.27%, palbociclib: 7.56%, no-boost DMSO = 9.72%, palbociclib = 13.39%.

*E,* Change in expression for E2F target pMHCs, plotted by source protein, with palbociclib treatment for the no-boost analysis and corresponding expression levels for the pMHC-boost analysis. X denotes pMHCs which were not quantified in the pMHC-boost analysis.

**Figure S3**

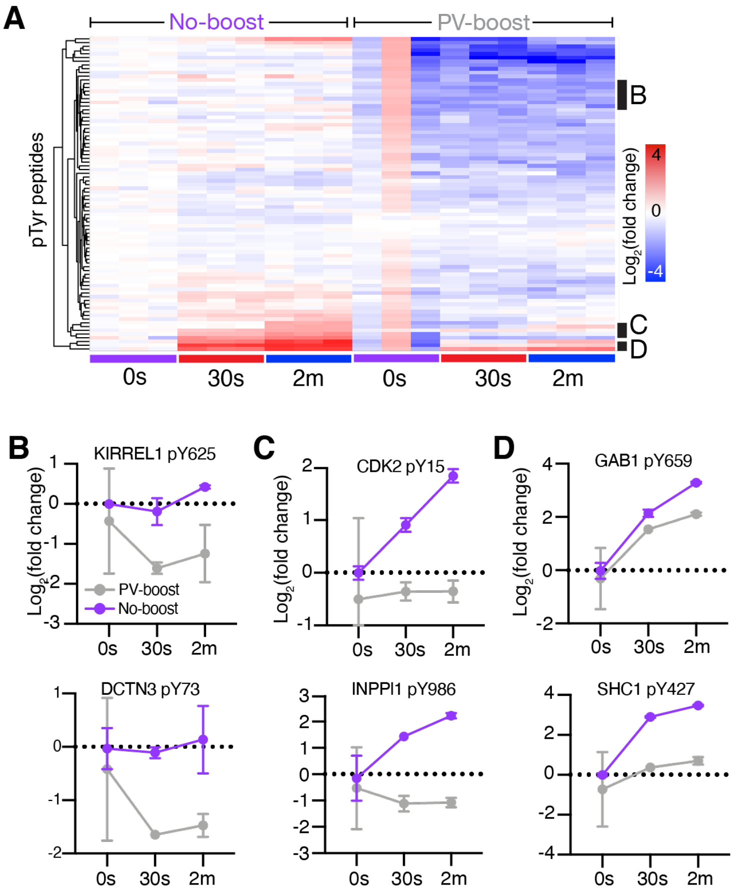


**Supplemental Fig. S3.**

*A,* Hierarchical clustering (Euclidean) of Log2(fold change) values of pTyr sites identified in both the PV-boost and no-boost analyses. Values are normalized to the mean 0s intensity.

*B-D,* Selected peptides from corresponding clusters highlighted in A represented as Log2(fold change) values over mean 0s intensities. Error bars represent +/- standard deviation.

**Figure S4**

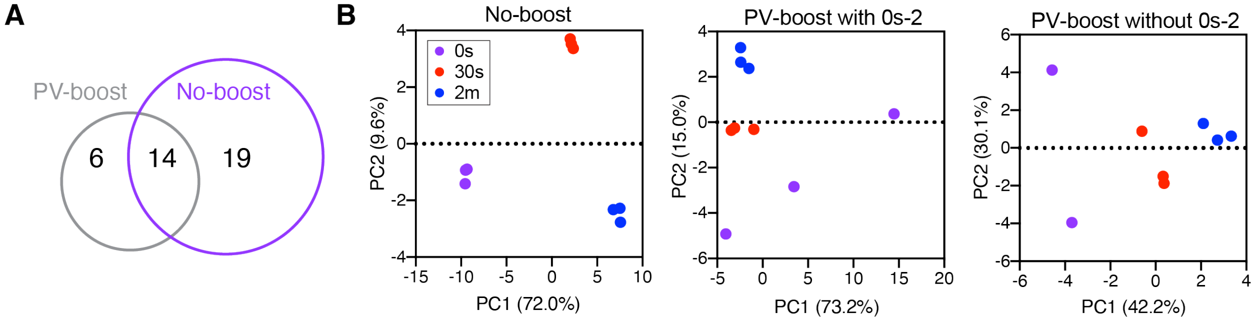


**Supplemental Fig. S4.**

*A,* Venn diagram of the number of unique pTyr peptides significantly different (two-tailed T test, $p<0.05$) between the 30s and 2m condition.

*B,* Samples plotted by principal component 1 (PC1) and PC2 score for no-boost and PV boost analysis including/excluding the 0s-2 sample and colored by EGF stimulation condition. Percentages define the variance described by the plotted PC.

**Figure S5**

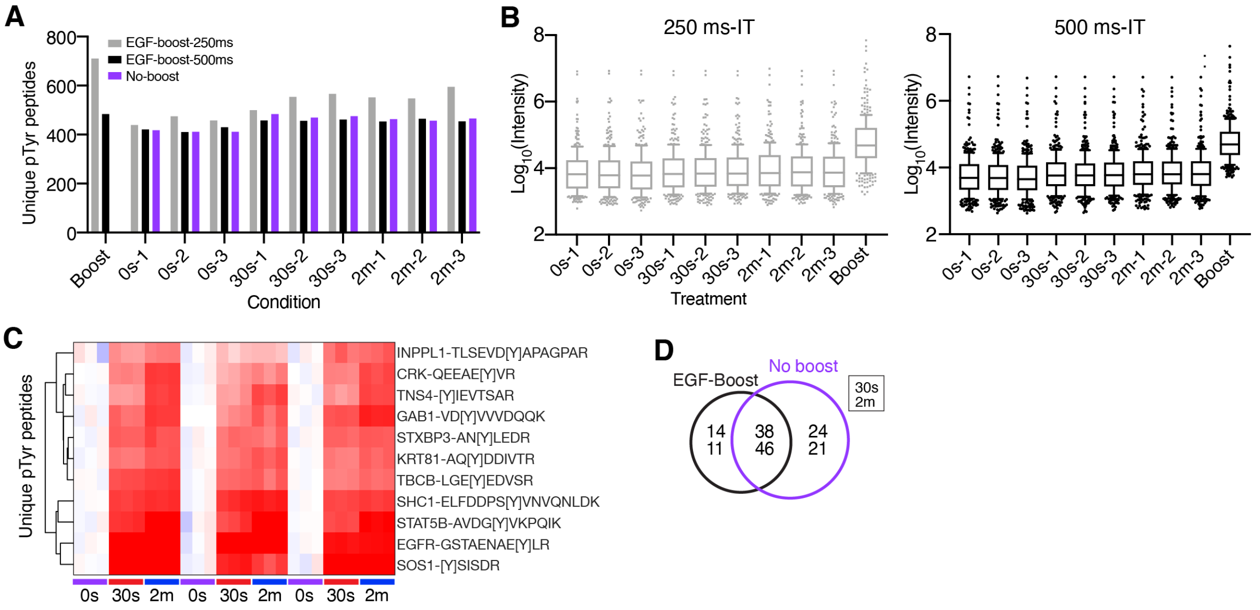


**Supplementary Fig. S5.**

*A,* Number of unique pTyr peptides quantified for each condition for each analysis.

*B,* Reporter ion intensities for EGF-boost 250 ms IT analysis. Boxes outline the interquartile range, and whiskers the 10 and 90th percentiles.

*C,* Hierarchical clustering of peptides highlighted by black bar in Figure 5A.

*D,* Venn diagram of number of pTyr peptides significantly different (two-tailed t-test, $p<0.05$) between the 0s and 30s (upper)/2m (lower) conditions using the EGF-boost 500ms and no-boost data.
